# Neuroprotection of low dose carbon monoxide in Parkinson’s disease models commensurate with the reduced risk of Parkinson’s among smokers

**DOI:** 10.1101/2023.05.27.542565

**Authors:** KN Rose, M Zorlu, X Xue, A Fassini, W Cai, S Lin, P Webb, MA Schwarzschild, X Chen, SN Gomperts

## Abstract

Paradoxically, cigarette smoking is associated with a reduced risk of Parkinson’s disease (PD). This led us to hypothesize that carbon monoxide (CO) levels, which are constitutively but modestly elevated in smokers, might contribute to neuroprotection. Using rodent models of PD based on α-synuclein (αSyn) accumulation and oxidative stress, we show that low-dose CO mitigates neurodegeneration and reduces αSyn pathology. Oral CO administration activated signaling cascades mediated by heme oxygenase-1 (HO-1), which have been implicated in limiting oxidative stress, and in promoting αSyn degradation, thereby conferring neuroprotection. Consistent with a neuroprotective effect of smoking, HO-1 levels in cerebrospinal fluid were higher in human smokers compared to nonsmokers. Moreover, in PD brain samples, HO-1 levels were higher in neurons without αSyn pathology. Thus, CO in rodent PD models reduces pathology and increases oxidative stress responses, phenocopying possible protective effects of smoking evident in PD patients. These data highlight the potential for low-dose CO modulated pathways to slow symptom onset and limit pathology in PD patients.

## INTRODUCTION

Parkinson’s disease (PD) is the second most common neurodegenerative disease, affecting an estimated 10 million individuals worldwide. While currently available therapies mitigate signs of disease, none of these limit neurodegeneration or alter the devastating course of the disease. Pathologically, PD is characterized by the formation and spread of alpha-synuclein (αSyn)-rich aggregates, called Lewy bodies, which have been proposed to induce degeneration of dopaminergic (DA) neurons in the substantia nigra pars compacta (SNpc) ^1^. Epidemiological research has identified smoking tobacco as the factor most consistently correlated with reduction in both the risk for developing PD ^2,3^, and the development of Lewy-body related neuropathology^4^. This finding prompted investigation into the therapeutic potential of tobacco smoke components, particularly nicotine. To date, however, nicotine has failed to mitigate PD symptoms or stall progression of disease in human clinical trials ^5,6^. Thus, other tobacco smoke constituents warrant evaluation.

One plausible, if paradoxical, candidate that has yet to be considered is carbon monoxide (CO). The concentration of hemoglobin-bound CO (CO-Hb) is intermittently higher in the blood of smokers compared to non-smokers, at levels usually below 10% that are well below those typically associated with the well-documented clinical or epidemiological toxicity of CO ^7–9^. Rather, at these levels, CO can activate cytoprotective signaling cascades mediated by Nrf2 and HIF-1α, which can reduce oxidative stress and inflammation ^10–17^. *In vitro,* low-dose CO also increases expression of Nurr-1, a transcription factor critical to the survival and maintenance of dopaminergic neurons ^18,19^. Notably, overexpression of heme oxygenase-1 (HO-1), a cytoprotective Nrf-2 and stress-induced enzyme that produces endogenous CO, has been found to protect dopaminergic neurons from neurotoxicity in an animal model of PD ^20^. Given the potential for CO to underlie the reduced risk of PD among smokers and to activate neuroprotective signaling, we hypothesized that low-dose CO might confer protection in rodent models of PD and that smokers might display higher levels of neuroprotective signaling factors.

## RESULTS

### Oral administration of low dose carbon monoxide reduces dopamine cell loss in an α-synuclein rat model of Parkinson’s disease

To investigate the neuroprotective potential of CO, we employed a genetic model of PD in which an AAV drives the overexpression of mutant A53T human αSyn in the SNpc of rats, resulting in the unilateral loss of dopaminergic neurons after 21 days ^21^. Injection of empty AAV1/2 vector into the contralateral SNpc provided a vector-matched within-animal control. Five days after AAV injection, rats received either CO in vehicle (HBI-002, Hillhurst Biopharmaceuticals) or the vehicle alone by enteral administration once daily for a total of sixteen days. Treatment with HBI-002 (10 ml/kg) elicited an increase in % CO-Hb peak to 4.5-7.8% (Figure 1a). A representative image showing the effect of CO on TH-positive neurons in the SNpc is shown in Figure 1b and quantitation is shown in Figure 1d. Administration of CO reduced ipsilateral loss of both TH-positive neurons in the SNpc (Figure 1b,d) and striatal dopamine (Figure 1e) when compared to rats treated with vehicle. To confirm that these effects were due to enhanced neuroprotection rather than a change in TH expression, we stained for the neuronal marker NeuN. A representative image showing the effect of CO on NeuN-positive neurons in the SNpc is shown in Figure 1c and quantitation is shown in Figure 1f. Compared to vehicle-treated animals, CO-treated animals had more NeuN-positive neurons in the ipsilateral SNpc (Figure 1c,f). Together, these data indicate that low-dose CO treatment increased survival of dopaminergic neurons in a progressive model of PD neurodegeneration.

**Figure 1.**
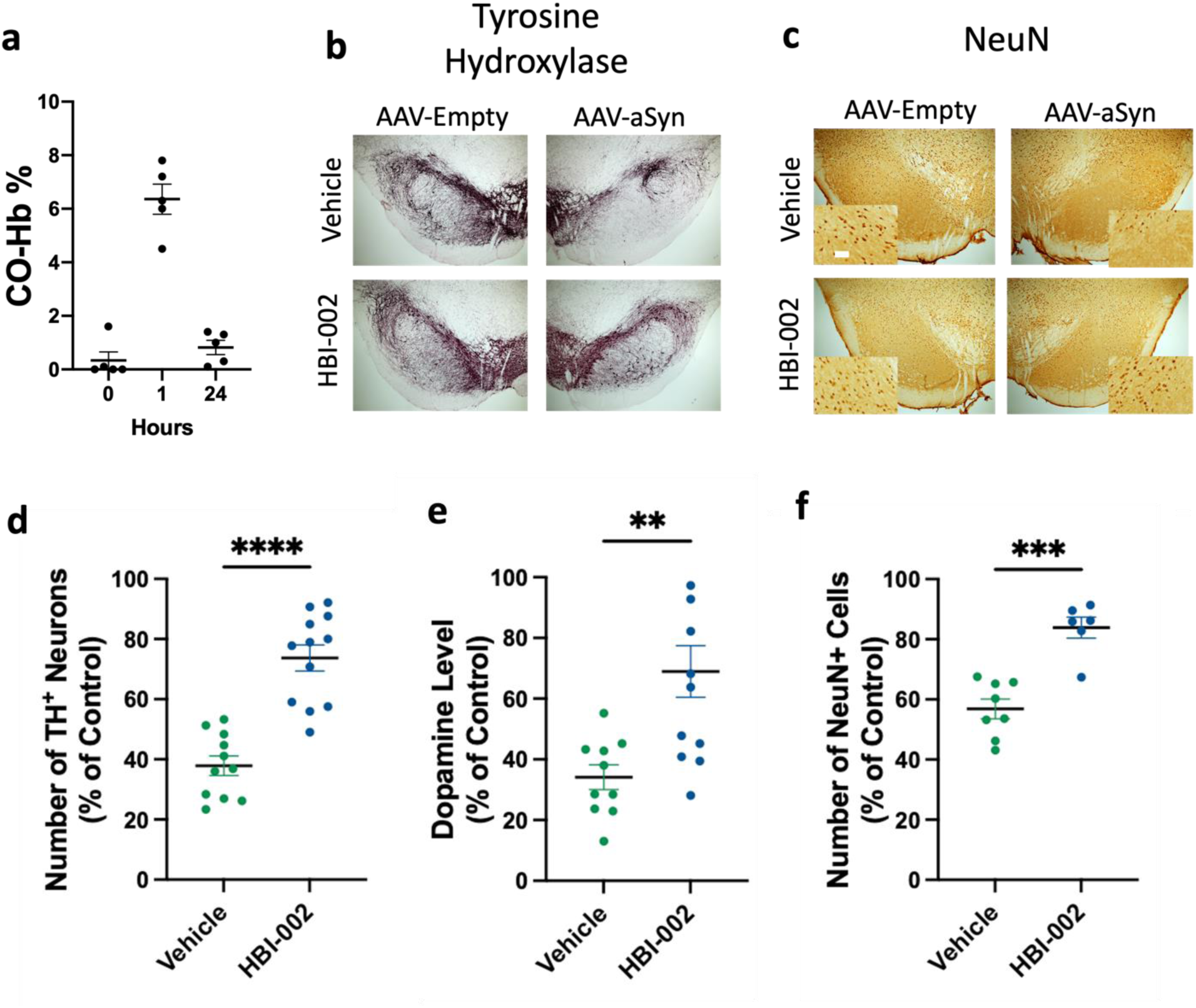
HBI-002, an oral formulation of CO, protects against human mutant A53T alpha-synuclein (aSyn) neurotoxicity. The left substantia nigra (SN) received AAV1/2-CMV-empty vector-WPRE-BGH-polyA (Control); the right SN received AAV1/2-CMV-human-A53T-alpha-sunuclein-WPRE-BGH-polyA. (**a**) Pharmacokinetics of CO-Hb% in blood before, 1 hour after, and 24 hours after a dose of HBI-002 (10mL/kg). **(b)** Representative photomicrographs of tyrosine hydroxylase (TH) in the SN. **(c)** Representative photomicrographs of NeuN in the SN. Scale bar, 20 μm. **(d)** Quantification of TH+ cells in the SN. Vehicle N=10, HBI-002 N=13, T-test, 1-β=0.99, P=0.0001. **(e)** Quantification of striatal dopamine by HPLC-ECD. Vehicle N=10, HBI-002 N=12, T-test, 1-β=0.92, P=0.0024. **(f)** Quantification of NeuN+ cells in the SN. Vehicle N=8, HBI-002 N=6, T-test, 1-β=0.99, P=0.001. Error bars represent the mean ± SEM.

### The protective effects of low dose carbon monoxide in the α-synuclein model are associated with increases in cathepsin D and reduced α-synuclein pathology

To determine the mechanism by which CO promoted survival of dopaminergic neurons, we next assessed whether CO administration affected cellular mechanisms of protein degradation, including autophagy and lysosomal function, as well as levels of αSyn and αSyn aggregates, which have been proposed to mediate neuropathology in PD patients ^1^. Markers of autophagy assessed by immunostaining of the ventral midbrain against LC3bI/II ratio, p62, Rab4, and Vsp35 remained unchanged with CO administration (Figure 2 a,b). By contrast, whereas no significant effect of CO treatment was evident on levels of the lysosome marker LAMP1 (Figure 2 a,b), levels of cathepsin D, an aspartyl protease found in the lysosome, were markedly elevated in ventral midbrains from animals treated with CO (Figure 2 a,b). As cathepsin D is one of the proteases responsible for αSyn degradation and its upregulation has been shown to be neuroprotective in PD models ^22–26^, we next evaluated the effect of CO on cathepsin D activity, as measured by cleavage of the cathepsin D substrate pepstatin A in differentiated SH-SY5Y human neuroblastoma cells. As shown in Figure 2 c,d, CO treatment significantly increased peptidase activity against a cathepsin D substrate.

**Figure 2.**
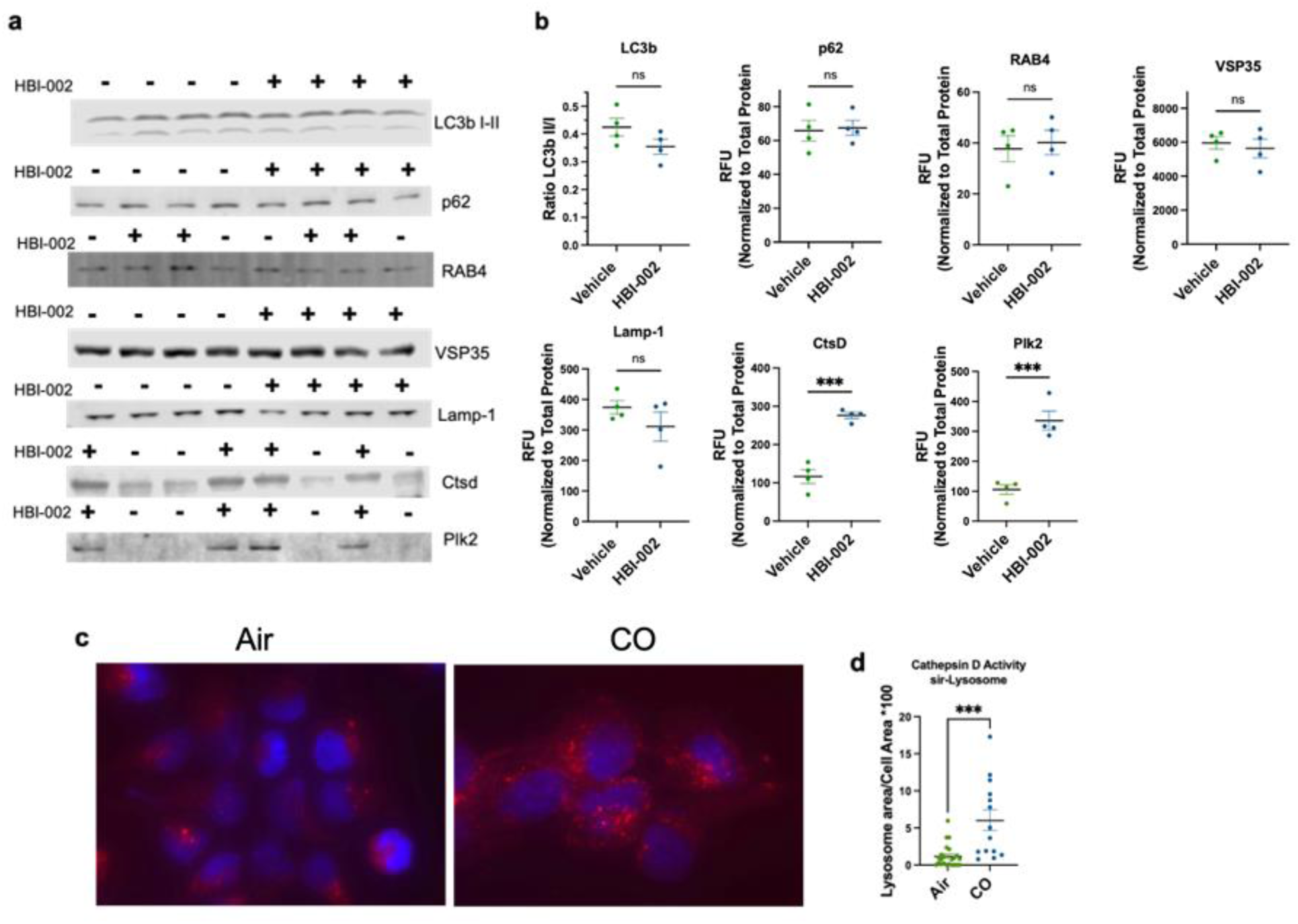
HBI-002 effects on autophagy, lysosomes and cathepsin D. **(a)** Immunoblots of autophagy and lysosomal activity markers. **(b)** Quantification of autophagy and lysosomal activity markers. Ctsd (p=0.002, 1-β=0.99, N=8, 4 per group) and Plk2 (p=0.006, 1-β=0.99, N=8, 4 per group), all other markers p>0.05. **(c)** Representative photomicrographs of sir-lysosome staining in SH-SY5Y human neuroblastomas. **(d)** Quantification of sir-lysosome staining p=0.0002, 1-β=0.95, N=38 cells, air treated n=23, CO-treated n=14. Error bars represent the mean ± SEM.

Because αSyn is a substrate of cathepsin D, and also, in its aggregated form, a principal pathological feature of PD ^27^, we next investigated whether low-dose CO might affect the levels of αSyn directly, and the levels of αSyn phosphorylated on Serine 129 (αSyn^pSer129^), a form of the protein associated with aggregates in PD patients, and also a pathological feature of PD-associated αSyn^A53T^ neurotoxicity in our rodent model ^28^. To do this, rats were subjected to the AAV and low-dose oral CO treatment protocol described above to induce ipsilateral but not contralateral PD-like neuropathology. After 3 weeks, tissue blocks containing the SN were harvested and subjected to western analysis (Figure 3) with αSyn or phospho-129-specific αSyn mAbs. αSyn mAb labeled several bands including a 14 KDa band consistent with αSyn monomer, while αSyn^pSer129^ labeled a high molecular weight band (260 KDa), suggesting that αSyn^pSer129^ is part of a homo- or hetero-protein aggregate (Figure 3a, Supplementary Figure 1). Whereas levels of αSyn monomer were similar between vehicle and CO-treated animals (Figure 3 a,b), levels of αSyn and αSyn^pSer129^ in the 260 KDa bands were significantly reduced in the CO-treated animals compared to vehicle controls (Figure 3 a,c,d). Thus, CO treatment reduced levels of a form of αSyn associated with aggregation and with neuropathology in PD patients and in our animal model ^29^.

**Figure 3.**
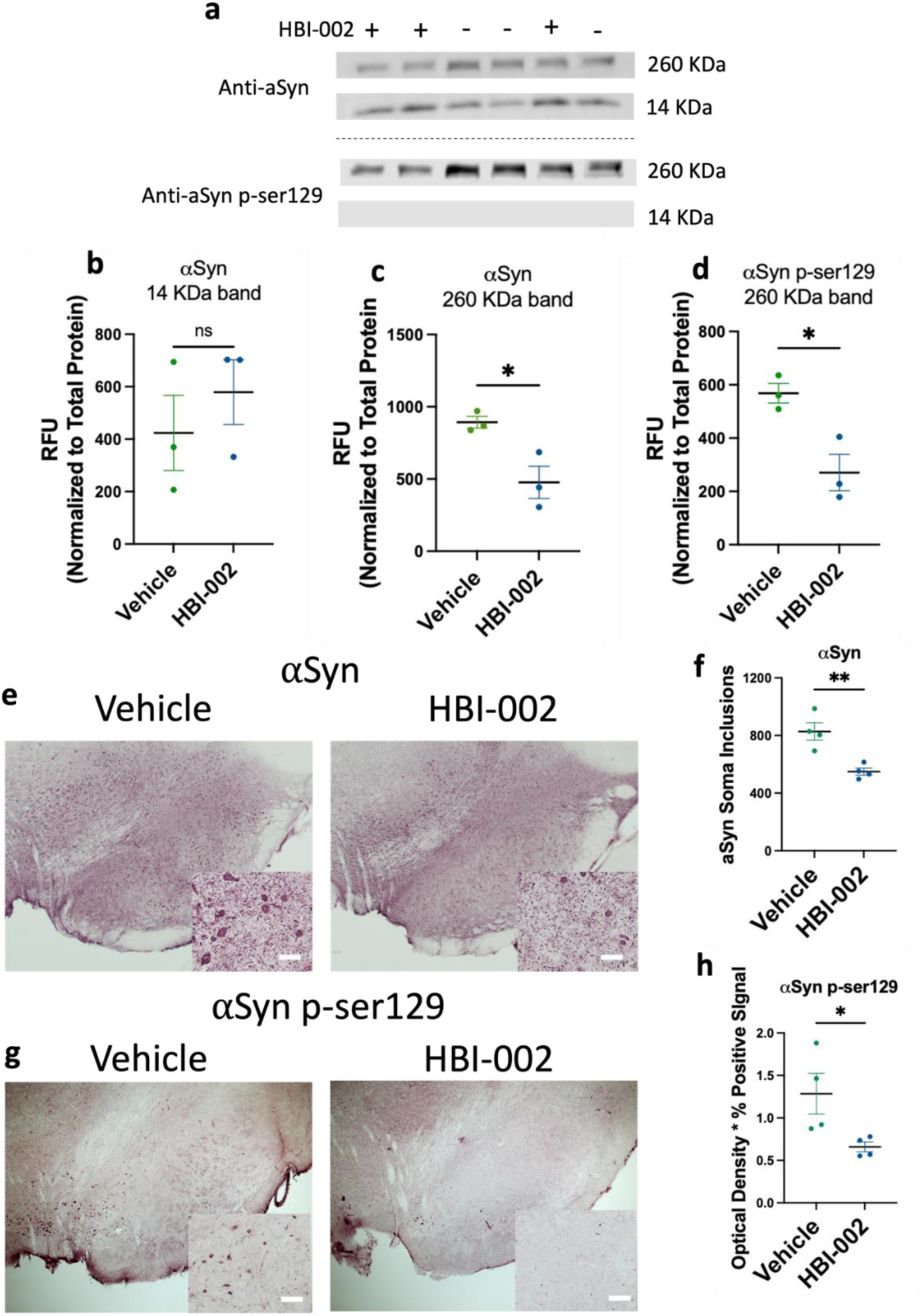
HBI-002 modulates αSyn pathology in an AAV-A53T model of PD. **(a)** Immunoblots of αSyn and pSer129-αSyn (260 KDa and 14 KDa). **(b)** Quantification of 14 KDa band of αSyn. P=0.69, 1-β=0.13, N=6, 3 per group. **(c)** Quantification of 260 KDa band of pSer129-αSyn. P=0.02, 1-β=0.88, N=6, 3 per group. **(d)** Quantification of 260 KDa band of pSer129-αSyn. P=0.0014, 1-β=0.81, N=6, three per group. **(e)** Representative photomicrographs of αSyn immunoreactivity in the SN. Scale bar, 20 μm. **(f)** Quantification of aSyn+ cells in the SN, values reflect the number of counted cells. P=0.02, 1-β=0.94, N=8 (4 per group). **(g)** Representative photomicrographs of pSer129-αSyn immunoreactivity in the SN. (**h**) Quantification of pSer129-aSyn. P=0.0014, 1-β=0.81, N=6, three per group. Error bars represent the mean ± SEM.

To test whether lesions associated with αSyn^pSer129^ aggregation in PD patients were affected by CO, we next interrogated serial sections of the SNpc from animals treated with oral CO or carrier by immunofluorescence with αSyn mAb or αSyn^pSer129^ mAb. Representative images are shown in Figure 3e,g and quantitation is shown in Figure 3f,h. In accordance with the western analysis, CO treatment reduced the overall immunoreactivity associated with αSyn mAb staining (Supplementary Figure 2). Further, as shown in Fig. 3e,f, CO treatment reduced the number of αSyn-containing immunoreactive puncta detected within the soma of neurons, a putative indicator of reduced αSyn aggregation. In line with the western analysis, CO treatment reduced the overall levels of immunoreactivity detected with αSyn^pSer129^ mAb (Figure 3 e,g). In accordance with these data, levels of Polo-like kinase 2 (Plk2), a serine-threonine kinase that phosphorylates αSyn at serine 129 (24) and facilitates degradation of αSyn (25, 26), were increased with oral CO treatment in the ventral midbrain in vivo (Figure 2a,b). Together, these data indicate that CO treatment reduced levels of a form of αSyn associated with aggregation and neuropathology in PD patients, and which contributes to neuropathology in the rat PD model.

### Low dose carbon monoxide reduces dopamine cell loss in an MPTP toxin model of Parkinson’s disease and increases the heme oxygenase-1 cytoprotective oxidative stress response pathway

To determine whether CO provided direct neuroprotection, we next evaluated its protective efficacy following administration of 1-methyl-4-phenyl-1,2,3,6-tetrahydropyridine (MPTP;40 mg/kg). MPTP induces rapid mitochondrial oxidative stress and proteasome disruption, and inflammation resulting in lesions within the nigrostriatal system ^30^, thus rapidly recapitulating many neurodegenerative features of PD evident in humans ^31^. To do this, male C57BL/6 mice received a single intraperitoneal dose of MPTP, and were then treated with either CO gas or with ambient air for an additional hour (see Methods); five days later, mice were sacrificed for analysis. Each treatment with CO gas increased CO-Hb to 15.6-23.2% (Figure 4a). Representative images are shown in Figure 4b and quantitation is shown in Figure 4c. In concordance with previous reports ^30^, MPTP induced significant loss of DA+ and TH+ neurons in the SN (Figure 4c,d); importantly, treatment with CO had no effect in saline treated animals but mitigated MPTP-induced toxicity in the nigrostriatal dopamine system (Fig. 4b,d). Consistent with these observations, CO had no effect on striatal dopamine in saline treated animals but reduced MPTP-induced loss of striatal dopamine (Figure 4c). Taken together, data from the αSyn^A35T^ model and MPTP model indicate that CO treatment can limit both αSyn S129 phosphorylation as well as neuronal degeneration associated with oxidative stress. Thus, we reasoned that the primary activity of CO might be to limit oxidative damage by inducing cytoprotective oxidative stress response pathways. To test this possibility, we set out to investigate CO effects on the cytoprotective protein HO-1 ^20^. HO-1 has been shown to be protective in animal models of PD, and its overexpression reduces αSyn aggregation ^32–34^. In addition, CO exposure reportedly increases HO-1 expression ^35–37^. We found that ventral midbrain HO-1 protein levels were significantly increased in rats treated with CO compared to vehicle controls (Figure 5a). To determine how CO upregulates HO-1 levels, we considered the possibility of CO-induced, HIF-1α mediated transactivation of HMOX1 ^38^. Indeed, ventral midbrains from rats treated with low-dose CO had significantly increased nuclear HIF-1α compared to those treated with vehicle (Figure 5 b,c), suggesting that CO-induced HIF-1α nuclear translocation may be responsible for the increase of HO-1.

**Figure 4.**
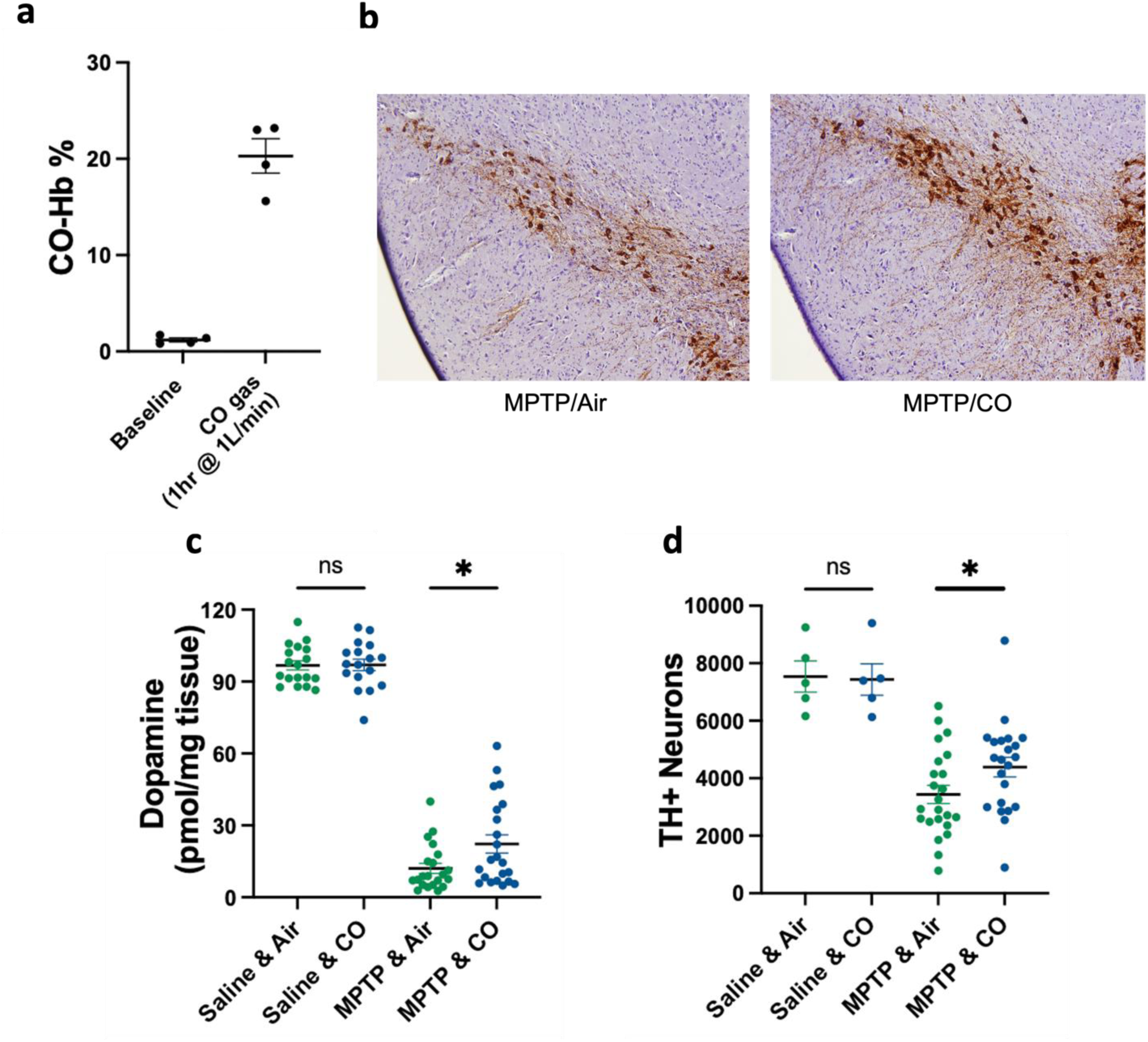
Carbon monoxide protects against MPTP induced neurotoxicity in the SN. (**a**) Concentration of CO-Hb in blood at baseline and after 1 hour of CO gas exposure (200 ppm at 1L/min). **(b)** Representative photomicrographs of TH immunoreactivity in the SN, counterstained with hematoxylin QS. **(c)** HPLC-ECD quantification of striatal dopamine content. **(d)** Quantification of TH-positive cells in the SNpc. Number of animals: Saline & air=5, Saline & CO=5, MPTP & air n=23, MPTP & CO n=22, P=0.04, 1-β=0.50. Number of animals: Saline & air n=18, Saline & CO n= 17, MPTP & air n=21, MPTP & CO n=22. P=<0.0001. Error bars represent the mean ± SEM.

**Figure 5.**
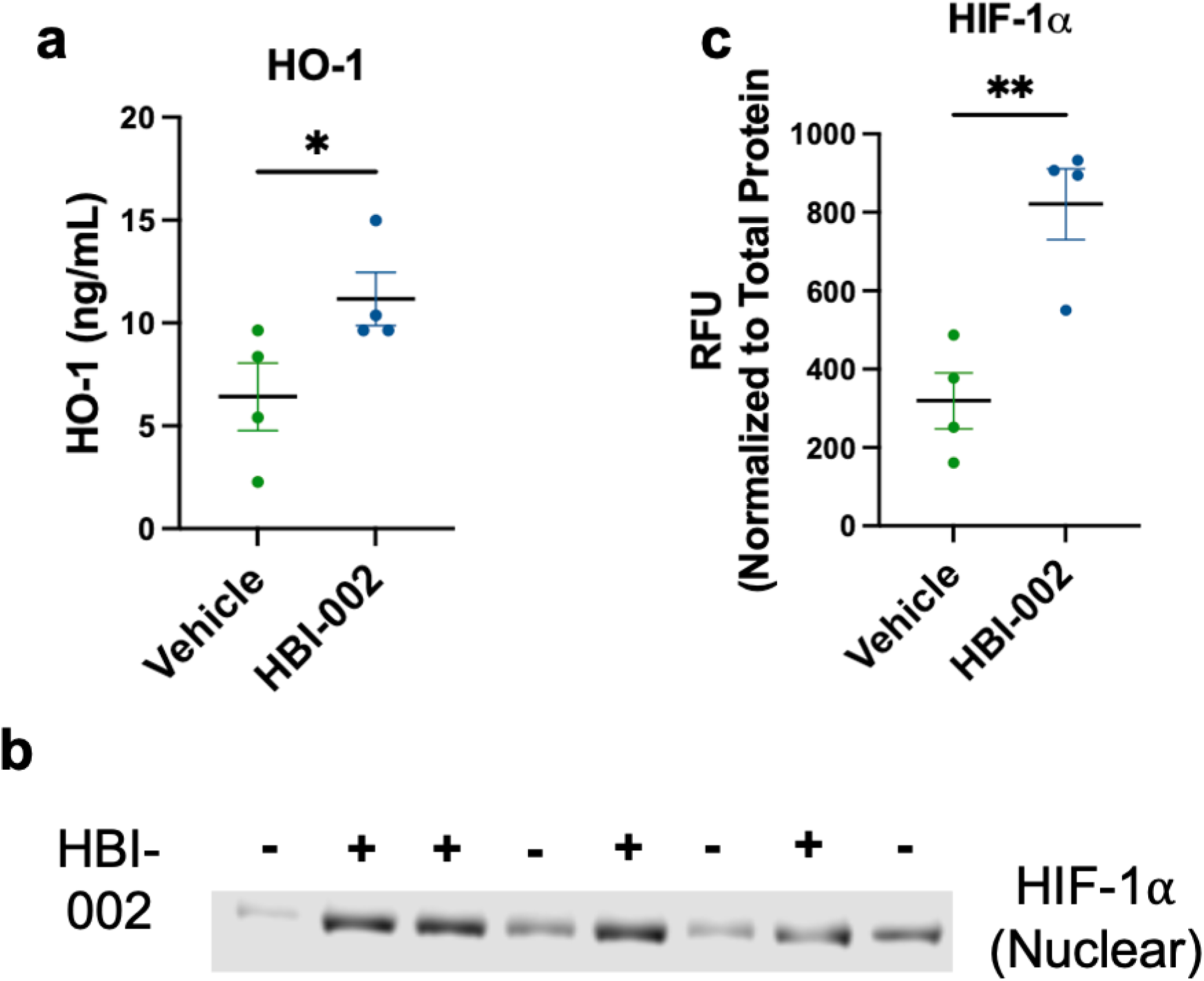
Carbon monoxide increases HO-1 in association with increased HIF-1α signaling. **(a)** Quantification of HO-1 by ELISA, P=0.0428, 1-β=0.47, N=8 (4 per group). **(b)** Immunoblots of nuclear HIF-1α. **(c)** Quantification of nuclear HIF-1α P=0.0048, 1-β=0.94, N=8 (4 per group). Error bars represent the mean ± SEM.

### Exposure to low dose carbon monoxide was not associated with neurotoxicity

While chronic inhalation of CO at doses associated with CO-Hb levels < 14% has not been associated with clinical toxicity in humans^9,10^ and animals^7,39,40^ CO dosed to exceed 40% CO-Hb is highly neurotoxic ^41^. To assess whether treatment with HBI-002 was associated with neurotoxicity, we evaluated CO-susceptible neuronal populations in rats, including Purkinje cell loss in the cerebellum and neuron degeneration in the cortex ^41–43^. As shown in Supplementary Figure 3a, administration of HBI-002 (10 mL/kg/day for 16 days) had no effect on the number of cerebellar empty basket cell counts (vehicle=59.8 ± 4.8 empty baskets, n=5, HBI-002=44.8 ± 10.8 empty baskets, n=6, p=0.27) ^44^. HBI-002 reduced the number of Fluoro-Jade C positive cells in cortex (Supplementary Figure 3b) (vehicle=12.7 ± 3.9, n=7, HBI-002=8.8 ± 2.8, n=7, p= 0.041). Thus, treatment with low-dose CO using HBI-002 was not associated with neurotoxicity.

### Human cigarette smoking is associated with increased CSF levels of heme oxygenase-1

Given the effects of low dose CO on cytoprotective cascades, we next sought to evaluate whether smoking is associated with neuroprotective markers in humans without PD. For this purpose, we examined HO-1 levels in cerebrospinal fluid (CSF) samples of smokers and nonsmokers. The demographic and clinical features of participants are shown in Table 1. The average ages of smokers (55.5 ± 15.6 years) and nonsmokers (54.3 ± 11.5 years) were comparable, as were the percentages of males across the groups (26% for smokers, 33% for nonsmokers). As shown in Figure 6a, HO-1 levels were elevated in smokers compared to nonsmokers. HO-1 levels did not vary with age (r = -0.03, p=0.8) or smoking intensity (packs per day; r = -0.05, p=0.7).

**Figure 6.**
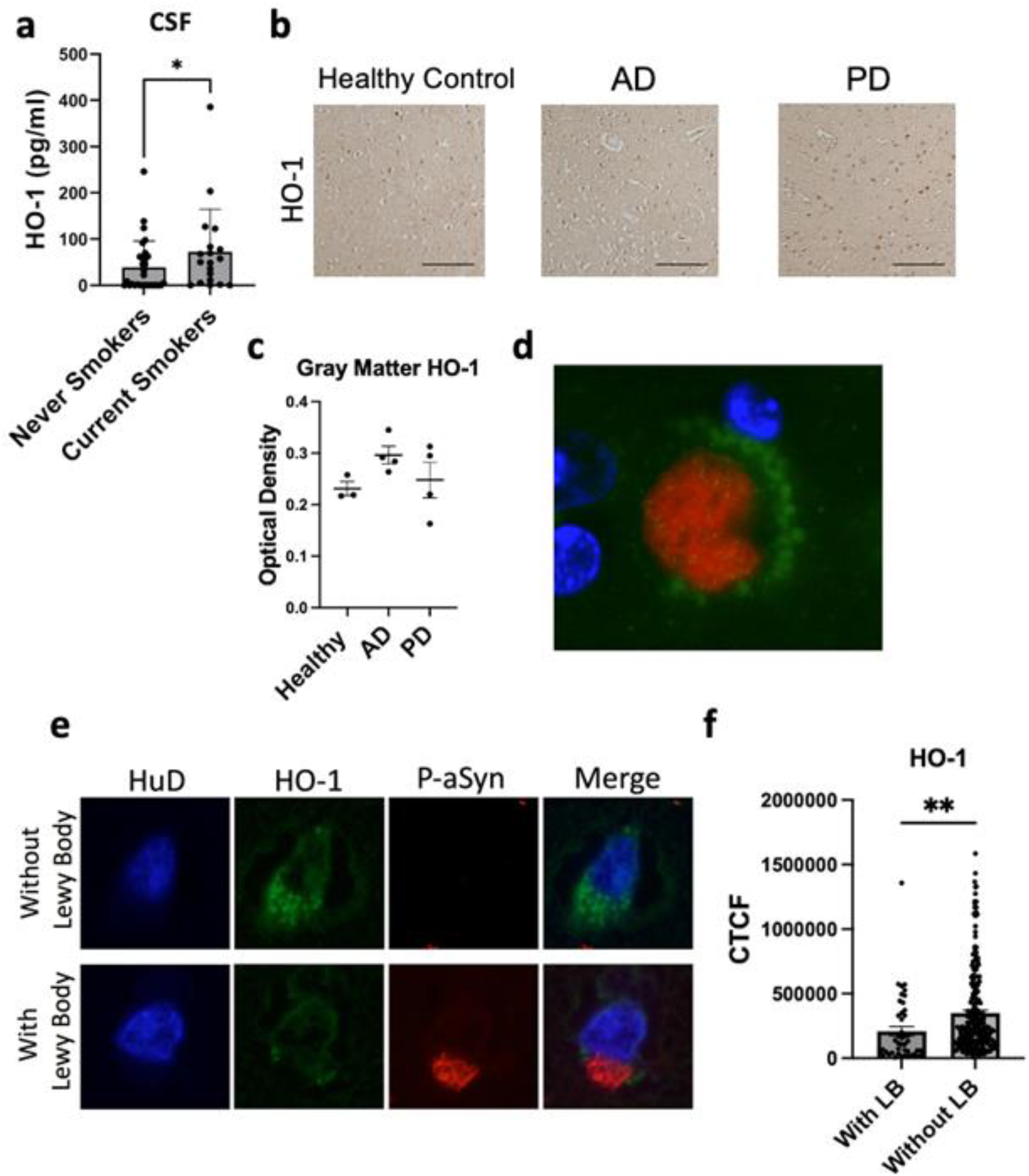
HO-1 in human samples. (**a**) HO-1 levels in cerebrospinal fluid (CSF) samples of smokers and nonsmokers without neurodegenerative disease, Wilcoxon one-tailed *p=0.040, N=49 (30 never smokers, 19 current smokers). **(b)** Representative photomicrographs of HO-1 immunoreactivity in the anterior cingulate of unaffected controls or patients with AD or PD. Scale bar, 50 μm. **(c)** Quantification of HO-1 immunoreactivity in gray matter of the anterior cingulate, p>0.05, 1-β=0.10 (3 healthy patients, 4 AD patients, 4 PD patients). **(d)** Illustrative photomicrograph of HO-1 (green) in close association with a Lewy body (p-ser129 alpha-synuclein, red) with DAPI nuclear counterstain (blue). **(e)** Representative photomicrographs of HO-1 in cells with and without Lewy bodies. Cells were co-stained for the neuronal-specific marker HuD (pseudocolored blue). **(f)** Quantification of HO-1 in PD samples in HuD-positive cells with and without Lewy bodies, **two-tailed p=0.0002, 1-β=0.83, N=279 HuD+ cells (46 with Lewy bodies, 233 without Lewy bodies) across 4 patients. CTCF: corrected total cell fluorescence (arbitrary units)

**Table 1.** Demographic characteristics of study participants providing CSF samples.

|  | Never smokers | Current Smokers | p value* |
| --- | --- | --- | --- |
| Age (mean, std) | 54.3 (11.7) | 55.5 (15.6) | 0.78 |
| Male (%) | 33.3% (10/30) | 26.3% (5/19) | 0.60 |
| **Clinical condition** |  |  |  |
| Idiopathic intracranial hypertension (IIH) | 33.3% (10/30) | 10.5% (2/19) | 0.09 |
| Normal pressure hydrocephalus | 13.3% (4/30) | 21.1% (4/19) | 0.69 |
| Headache (non-IIH) | 20% (6/30) | 26.3% (5/19) | 0.73 |
| Neuropathy, radiculopathy | 13.3% (4/30) | 10.5% (2/19) | 1 |
| Psychiatric** | 3.3 (1/30) | 21.1% (4/19) | 0.07 |
| Other*** | 16.7 (5/30) | 10.5% (2/19) | 0.69 |
| Packs-per-day (mean, std) | - | 0.48, 0.37 | - |

\*Chi-square test was used for all contrasts except age, where Student's t-test was used.
\*\*Psychiatric clinical condition includes participants with schizophrenia, psychosis, anxiety, and functional neurologic disorder.
\*\*\*Other includes a healthy volunteer and participants with no neurologic disease, neck pain, multifactorial gait disorder, isolated subjective cognitive complaints, spells of uncertain etiology, and sleep disorder.

### Heme oxygenase-1 in human brain tissue

To determine whether HO-1 levels are altered in human PD brain tissue, we evaluated HO-1 in anterior cingulate cortex samples from PD patients harboring neuropathologically proven Lewy body disease, Alzheimer’s disease (AD), or age-matched normal controls. Overall, HO-1 levels were comparable across PD, AD, and control tissue (Figure 6b,c). These findings contrast with the previously reported increase in HO-1 in PD individuals in the substantia nigra ^45^ and in saliva ^46^. However, when we co-stained for the Lewy body marker αSyn^pSer129^ and HO-1, we observed a close association of HO-1 puncta surrounding many Lewy bodies (Figure 6d). Thus, the association of HO-1 with Lewy bodies is not unique to PD dopamine cells ^47^. Even so, we observed that neurons harboring Lewy bodies had significantly lower levels of HO-1 when compared to adjacent neurons lacking Lewy bodies (Figure 6e,f). Together with the effects of low-dose CO on reducing TH+ cell loss, on reducing aggregated and phosphorylated species of αSyn, and on increasing HO-1, these neuropathological observations in human PD brain tissue suggest that neuronal HO-1 in PD may reflect a potentially protective response to PD-associated neuronal injury.

## DISCUSSION

Smoking is associated with a reduced risk of PD ^2,3^, but the mechanisms underlying smoking mediated neuroprotection have been unclear. In this study, we tested the hypothesis that low doses of carbon monoxide (CO), which are constitutively and intermittently elevated in individuals who smoke, might contribute to neuroprotection. We demonstrate in genetic and toxin models of PD that treatment with low-dose CO preserves SNpc dopamine neurons and associated striatal dopamine. Further, we show that CO treatment reduces the accumulation of αSyn within neuron somas and reduces the phosphorylation of αSyn that is associated with toxicity. These results suggest that administration of low-dose CO in rodent PD models may phenocopy neuroprotective effects of smoking seen in PD patients.

While there are numerous contaminants in tobacco smoke, CO is a major one. Nevertheless, the hypothesis that CO is associated with neuroprotection in PD is counter intuitive, as CO is thoroughly well-documented as a neurotoxin at high concentration and is generally considered an environmental toxin. It is also well-documented that peak CO-Hb levels in smokers are almost always below 15% and fluctuate given the episodic nature of the habit. Clearly documented adverse CO-related effects at 15% CO-Hb have not been identified other than limiting extensive exercise, some individuals with myocardial ischemia as well as with the developing fetus^7^. Further, multiple Phase 1 and Phase 2 clinical studies in both healthy and disease impacted individuals have failed to identify clinically relevant adverse effects of CO at peak CO-Hb levels below 15% (clin trials.gov and ^9^). As all drugs cause toxicity at sufficient dose and other well-documented toxins have found important clinical applications, it will be important to understand the risk-benefit of CO at low levels in patient populations such as PD.

The efficacy of low-dose CO in PD models draws attention to the PD-relevant pathways modified by CO. Consistent with prior reports implicating failure of lysosomal activity and cytoprotective cascades in PD ^48,49^, CO’s neuroprotective effects were associated with upregulation of cathepsin D, Plk2, HIF-1α, and HO-1. The engagement of multiple PD-relevant mechanisms is consistent with the reported pleiotropic activity of CO, which binds with high affinity to heme containing proteins in the body and the brain ^50^. Cathepsin D not only increases lysosomal clearance of αSyn, but also generates saposin C, a cofactor required for glucocerebrosidase function that has been implicated in PD ^51^. PLK2 also regulates αSyn clearance, reducing its toxicity ^52–54^. In addition, CO’s induction of HO-1 is likely also disease-relevant, as overexpression of HO-1 is protective in PD models ^20,32,55^, polymorphisms in HMOX1 have been associated with PD ^56^, and increased HO-1 levels have been reported in saliva samples in early PD ^57^.

In activating lysosomal and cytoprotective cascades, CO exposure recapitulates effects observed with cigarette smoke exposure. In mice, inhalation of cigarette smoke has been shown to increase peripheral cathepsin D expression and activity ^58^ as well as HIF-1α and HO-1 expression ^59,60^. Consistent with these observations, human smokers also have increased peripheral cathepsin D expression and activity ^61^, and we observed modestly higher HO-1 levels in the CSF of smokers compared to never smokers. Together, these results support the hypothesis that CO exposure contributes to smoking’s activation of these lysosomal and cytoprotective cascades and potentially thereby to the reduced risk of PD among smokers.

Building on the prior observation in PD brain tissue that HO-1 can be observed in Lewy body-laden dopamine cells, we found that HO-1 could be identified in Lewy body-containing neurons in the anterior cingulate cortex as well, another brain region affected early in PD. Even so, HO-1 levels were higher in neurons lacking Lewy bodies than in neurons harboring them. Together with evidence of CO-mediated neuroprotection in PD models, this observation raises the possibility that neuronal HO-1 upregulation may reflect a potentially cytoprotective, compensatory response to PD-associated injury. Further experiments will be required to test this hypothesis.

A limitation of the current study is the use of a viral vector to induce αSyn overexpression for model generation. While no proven translational model of PD exists, AAV provides a rapid and reproducible model that recapitulates many aspects of the disease, including αSyn accumulation and Lewy body-like somatic accumulation reminiscent of human cases of PD ^21^. However, αSyn overexpression is a feature rarely seen in cases of sporadic PD and is thought to be present only in cases with αSyn gene duplication or triplication ^62^. Furthermore, overexpression limits the ability to dissect mechanisms of αSyn degradation. We demonstrated through various methods and multiple cohorts of animals that CO treatment reduces accumulated and phosphorylated αSyn. However, it is unclear if the upregulation of degradation pathways acts on monomers, oligomers, or heavy-weight accumulations. Although there were no statistical differences in monomer expression, monomer overexpression may not allow for thorough investigation of the effect of CO on monomer degradation. Using longer-term PD models that do not require overexpression such as pre-formed fibril injections to striatum or peripheral nerves may be helpful to better understand CO’s effect on αSyn degradation. Additionally, causal studies using genetic or pharmacological knockdown of cathepsin D, PLK2, HIF-1α, and HO-1 would further implicate these molecules as neuroprotective in PD.

As expected based on prior CO toxicology studies ^7,39,40^, including chronic CO dosing studies of 3 to 24 months duration with peak CO-Hb 5 to 20% in multiple species, including mice, rats, dogs, guinea pigs, and monkeys, reporting no toxicological findings ^39,40^, we observed no evidence of CO-associated neurotoxicity at the low doses studied here.

Low-dose CO, as defined by levels associated with minimal toxicity, has recently been demonstrated to confer protection in numerous diseases, including traumatic brain injury, stroke ^11,12^, sickle cell disease ^63^, and colitis ^64^. Together with the well-established reduced risk of PD among smokers ^2,3^, the neuroprotective effects of low-dose CO demonstrated in both an αSyn genetic model and an MPTP toxin model of PD, the engagement of PD-relevant cascades, and findings in smokers and in human PD brain tissue that support HO-1 engagement increase the translational potential of low-dose CO for PD. Whether low-dose CO is safe in patients with PD has yet to be determined but further investigation appears warranted, given reports that PD smokers have tended to have fewer deaths from neurologic causes than PD nonsmokers, with no significant association between smoking status and all-cause mortality ^65^. Until low-dose CO is proven safe in PD, low-dose CO will nonetheless have value as a modulator of PD-relevant pathways with utility for its dissection and treatment.

In the context of the epidemiologic foundation that has identified smoking tobacco as a large inverse risk factor for PD ^2,3^, these results support the potential of pathways modified by low-dose CO to slow disease progression in PD. The present results demonstrating neuroprotection in PD models support further investigation of these pathways in PD.

## MATERIALS AND METHODS

### Study Design

The overall goal of this work was to evaluate the potential of low-dose CO to engage PD-relevant cascades and reverse neuropathology in models of PD. For this purpose, we studied an oral liquid drug product containing CO currently under development, HBI-002 (Hillhurst Biopharmaceuticals), and CO gas. We studied the efficacy of HBI-002 (oral gavage, 10 ml/kg) compared to HBI-002 vehicle in the AAV-aSynA53T rat model of PD (n = 13 per group). We studied the efficacy of CO gas (inhaled, 200 ppm) compared to air in the short term MPTP model of PD (n = 40 per group). Investigators were blinded to the HBI-002 versus HBI-002 vehicle intervention and were blinded for all analyses. Blinding was achieved by using non-descript, random code letter combinations for samples. Blinding was broken only after data had been collected and input into statistical programs. Injection location was verified in every animal by visual inspection of the wound tract and α-synuclein expression in target regions. Two animals were removed from the analysis because the stereotactic injection missed the substantia nigra. Another animal was excluded from dopamine analysis as dopamine levels were well above 100% of control, possibly due to improper dissection of the striatum. This animal was identified as an outlier by the ROUT method. Each study was replicated 2 times. The data provided are biological replicates of a single study.

### Animals

Six-month-old male C57BL/6 mice were purchased from The Jackson Laboratory (Bar Harbor, ME), and female Sprague Dawley rats ranging from 220-240 g were purchased from Charles River (Wilmington, MA). All animals were housed in the Center for Comparative Medicine at Massachusetts General Hospital’s Institute for Neurodegenerative diseases with a 12hr light/dark cycle and access to food and water *ad libitum*. All experiments were approved by Massachusetts General Hospital’s institutional animal care and use committee (protocol # 2015N00014, 2019N00168).

### Overexpression of human A53T α-synuclein in the SNpc of rats

Rats were anesthetized with isoflurane/oxygen and underwent bilateral stereotactic surgery. Each animal received 2 μL of a 5×10^12^ GC/mL viral titer injected into the nigra at the following coordinates: AP: -5.2, ML: +/- 2.0, DV: -7.8. The left nigra received AAV1/2-CMV-empty vector-WPRE-BGH-polyA; the right nigra received AAV1/2-CMV-human-A53T-α-synuclein-WPRE-BGH-polyA. Viruses were purchased from Vigene Biosciences (Rockville, MD). Animals were allowed to recover for 5 days before interventions began.

### Oral carbon monoxide treatment

A proprietary liquid carbon monoxide drug product currently under development, referred to as HBI-002, was provided by Hillhurst Biopharmaceuticals, Inc. (Montrose, CA). HBI-002 or vehicle (HBI-002 vehicle without CO) was given to animals via oral gavage (14 GA needle) at a dose of 10 mL/kg/day. Animals displayed no signs of distress and had no appreciable change in serum lactate in response to carbon monoxide dosing (data not shown).

### HBI-002 and inhaled gaseous CO pharmacokinetics

After a single maximum feasible dose of CO (limited by stomach volume) given by oral gavage or gaseous administration, animals were deeply anesthetized with an overdose of ketamine/xylazine followed by cardiac puncture into the right ventricle. Blood was collected using a 3 mL heparin coated line-draw syringe and a 23 Ga. needle. Blood samples were then immediately analyzed for CO-Hb% by oximetry using an Avoximeter 4000 (Instrumentation Laboratory, Bedford, MA). In rats, a 10 ml/kg dose of HBI-002 elicits an increase in % CO-Hb peak to 7.6-8.9%, whereas in mice, the same dose increases % CO-Hb to 2.6-3.1% (Supplementary Figure 4). In contrast, treatment with CO gas (200 ppm) increased % CO-Hb in mice to 19.4-23.3%.

### MPTP administration in mice and gaseous CO administration

Mice were given an i.p. injection of 40mg/kg MPTP or saline. One hour following injections, mice were treated with CO gas (200 ppm, 1 L/min) or air (1 L/min) for one hour by placing animals in a chamber with a constant flow of gas. Five days after treatment, animals were sacrificed via cervical dislocation and their brains harvested for further processing. CO gas was employed for these experiments instead of HBI-002 as the pharmacokinetics of oral HBI-002 are less favorable in mice.

### Quantification of dopamine by HPLC-ECD

Animals were deeply anesthetized with ketamine and xylazine followed by rapid decapitation and brain removal. The striatum from each hemisphere was segregated, dissected on ice, and frozen on dry ice. Pieces of frozen striatum were weighed and homogenized in 0.1 mM EDTA, 1 μM 3,4 dihydroxybenzylamine hydrobromide (DHBA, internal standard), and 50 mM phosphoric acid in a 1:20 ratio (weight: volume). The resulting homogenate was centrifuged at 14,000 x g to pellet cell debris and precipitated protein. The supernatant was then filtered through Costar SpinX 0.22-micron spin filter cartridges (Sigma-Aldrich). After filtering, 5 μL of supernatant was injected onto a Microsorb-MV column (C18, 150 mm x 5.6 mm, 5 micron) using an Ultimate 3000 UHPLC system (Thermo Fisher). Separation was achieved with a 17-minute isocratic method at a flow rate of 0.6 mL/min, and a mobile phase consisting of 75 mM sodium phosphate monobasic, 1.75 mM sodium-1-octanesulfonate, 100 μL/L triethylamine, 25 μM EDTA, and 10% acetonitrile. Detection was carried out with an Ultimate 3000 ECD-3000RS (Thermo Fisher) with a screening electrode set to -150 mV and a detection electrode set to 250 mV. DHBA was used as an internal standard and dopamine concentration was calculated from a standard curve. Analyses were blinded to experimental conditions.

### Immunohistochemistry

Blocks of tissue containing the SNpc were drop-fixed in 4% paraformaldehyde for three days. After that time, brains were sectioned at 40 μm on a vibratome (Leica, Buffalo Grove, IL). Sections were collected in a 1 in 6 series and stored in PBS at 4°C. For each antigen, a single series of sections were stained from each animal. Free-floating sections were incubated in 3% hydrogen peroxide for 15 minutes to block endogenous peroxidase. Next, sections were blocked and permeabilized in PBS with 2.5% bovine serum albumin (BSA), 10% normal goat serum (NGS), and 0.3% Triton X-100 for 30 minutes. Sections were then transferred to wells containing the corresponding primary antibody diluted in PBS with 2.5% BSA and 10% NGS. Primary antibodies used were tyrosine hydroxylase (Abcam cat# ab112, 1:1000), NeuN (BioLegend cat# 834501, 1:2000), alpha-synuclein (ThermoFisher cat# 32-8100, 1:1000), p-ser129 alpha-synuclein (BioLegend cat# 825702, 1:1000), calbindin D (ThermoFisher cat# MA5-50510, 1:1000), and glutamic acid decarboxylase (GAD) (ThermoFisher Cat# MA5-24909, 1:100). Sections were allowed to incubate overnight at 4°C. After several washes in PBS, antigens were visualized using an avidin-biotin detection system (ABC elite kit, Vector, Burlingame, CA) with ImmPact VIP and DAB substrates (Vector, Burlingame, CA) following the manufacturer’s instructions. Sections were mounted, dehydrated in graded ethanol, cleared in xylene, and cover slipped with permanent mounting media (VectaMount, Vector, Burlingame, CA). In instances where sections were counterstained, hematoxylin QS (Vector, Burlingame, CA) was used per the manufacturer’s instructions. For staining calbindin and GAD, sections underwent antigen retrieval by boiling sections in sodium 20 mM citrate buffer (pH=6) for 20 minutes before continuing with the staining procedure.

### Human CSF samples

Samples were obtained from the Mass General Institute for Neurodegenerative Disease (MIND) biorepository, which includes CSF collected with research use consent at the time of diagnostic lumbar punctures at the Department of Neurology at Massachusetts General Hospital (IRB: 2015P000221). De-identified samples were collected and aliquoted in polypropylene plasticware, labelled, frozen on dry ice within 30 min of collection, and stored at −80 C until use. Inclusion criteria for participant samples studied here included the absence of neurodegenerative, inflammatory, or immune mediated diseases, and the absence of inflammatory CSF. We further required that samples were negative for Alzheimer’s disease on the basis of ADmark or in house testing of CSF Ab42/40, pTau181, and tTau using Euroimmun ELISA assays (Lübeck, Germany) on a semiautomated Tecan Freedom Evo liquid handler (Männedorf, Switzerland). CSF samples from 30 never smokers and 19 active smokers were identified.

### Human tissue

Brain tissue samples from 4 donors with PD, 4 donors with AD, and 3 donors without neurodegenerative disease were selected from the Massachusetts Alzheimer’s Disease Research Center (MADRC) Neuropathology Core, which obtained written informed consent from next-of-kin for brain donation. Brains donated to the MADRC have undergone full brain autopsy and neuropathological confirmation of diagnosis, with comprehensive assessment of primary and co-pathologies. Brains from de-identified PD patients had high Lewy body pathology (Lewy body disease Braak stage 5-6) and met criteria for Lewy body disease ^66^. Brains from AD patients had high amyloid and tau pathology and no Lewy body disease pathology (Thal stage 4-5, Braak stage VI, LBD Braak stage 0) and met criteria for Alzheimer’s disease according to NIA-AA guidelines ^67^. Tissue from the anterior cingulate region was used. Tissue from the SNpc was not used due to severe neurodegeneration in late stages of the disease that leaves very few neurons and Lewy bodies for analysis.

For staining human tissue, formalin-fixed, paraffin embedded sections were cleared in xylene and brought to TBS through graded ethanol and water. Antigen retrieval was performed by boiling sections in 10mM citric acid (pH=6) for 30 minutes followed by a 20-minute cooldown. Lipofuscin was blocked using TrueBlack (Biotium, Freemont, CA) following the manufacturer’s instructions. Sections were then blocked with 10% BSA and 2.5% NGS. The primary antibodies used were p-ser129 alpha-synuclein (FujiFilm cat# 015-25191, 1:1000) and HO-1 (Cell Signaling cat# 26416, 1:200). Sections were allowed to incubate overnight at 4°C. Following several washes in TBS, sections were incubated in anti-mouse Alexa Fluor 555 and anti-rabbit Alexa Fluor 488 (Thermo Fisher cat# A-21422 and A-11008, respectively). After several washes in TBS, sections were incubated with anti-HuD directly conjugated to Alexa Fluor 647 (Santa Cruz cat# sc-28299, 1:50) for 1 hour at room temperature, followed by several washes in TBS, cover slipping with anti-fade mounting media, and imaging. ImageJ was used for staining intensity quantification. Analyses were blinded to experimental conditions when possible.

### Stereology

Sections labeled with TH, NeuN, and α-synuclein (Syn211) underwent counting using the optical fractionator principles with CAST stereology software (Olympus, Tokyo, Japan). Experimenters were blinded to experimental conditions. Counting was limited to the substantia nigra and was done with a 20X objective with a meander sampling of 100% to count every cell in the entire region. A total of six sections spanning the SNpc were counted per animal.

### ELISA

ELISA was used to quantify HO-1 levels in rat midbrain samples (Enzo Life Sciences cat#ADI-EKS-810A) and human CSF samples (Abcam cat#ab207621). HO-1 concentration was determined using a standard curve, and values are reported as ng HO-1/mg protein for rat samples and pg HO-1/ml of human CSF. Analyses were blinded to experimental conditions.

### Protein Extraction

Using a matrix, 3 mm blocks of brain tissue containing the SNpc were dissected. Then, a 3 mm biopsy punch was used to dissect the tissue containing the SNpc. The tissue punches were then homogenized in TBS with a protease and phosphatase inhibitor cocktail (Sigma Aldrich, St Louis, MO). The resulting homogenate was cleared by ultracentrifugation at 100,000 x g for 1 hour at 4°C. For nuclear protein extractions, fresh tissue was homogenized in Dounce homogenizers using NE-PER (Thermo Fisher cat#78833) following the manufacturer’s instructions.

### Immunoblotting

Samples were diluted in 2x Lamelli buffer and heated at 70°C for 10 minutes. Samples were then loaded on a 10-20% polyacrylamide gel and electrophoresis was carried out at 225 V until the dye front reached the bottom of the gel. Then, proteins were transferred to PVDF membranes, washed in distilled water, and dried overnight at 4°C. After reactivation of the PVDF membrane in methanol, total protein per lane was quantified using Revert 700 total protein stain and an Odyssey CLx imaging system (Licor, Lincoln, NE) on the 700 nm channel per the manufacturer’s instructions. Membranes were then blocked in 5% dry non-fat milk powder in TBS for 1 hour at room temperature followed by overnight incubation at 4°C in primary antibody diluted in 5% dry non-fat milk powder in TBS-T. Primary antibodies used were: alpha-synuclein (ThermoFisher cat# 32-8100, 1:1000), p-ser129 alpha-synuclein (FujiFilm cat# 015-25191, 1:1000), LC3b (Novus cat# NB600-1384, 1:1000), p62 (Abcam cat# ab109012, 1:10,000), RAB4 (Abcam cat# ab109009, 1:1000), VSP35 (Invitrogen cat# PA5-21898, 1:1000), Lamp-1 (Invitrogen cat# MA5-29384, 1:500) Ctsd (Cell Signaling cat# 69854, 1:1000), Plk2 (Novus cat# NBP2-15078, 1:500), Nrf2 (Novus NBP1-32822, 1:1000), and HIF-1α (Novus cat# NB100-449, 1:1000). After several washes in TBS-T, membranes were incubated in a 1:30:000 dilution of IRDye 800 donkey anti-rabbit or anti-mouse (Licor, Lincoln, NE) in 5% dry non-fat milk powder and 0.02% SDS in TBS-T for 1 hour at room temperature. For p-ser129 α-synuclein, 50mM NaF was added to all milk-based buffers. After several washes in TBS-T followed by a final wash in TBS, blots were imaged with an Odyssey CLx imaging system on the 800 nm channel. Fluorescent intensity of bands was determined using Image Studio (Licor, Lincoln, NE) and bands were normalized to total protein loaded in individual lanes. Three technical replicates were used for quantification purposes. Analyses were blinded to experimental conditions.

### Cell culture

Undifferentiated human neuroblastoma SH-SY5Y cells were purchased from American Type Culture Collection (ATCC, VA, USA) and maintained in high glucose DMEM supplemented with 10% FBS and 1% P/S. Cells were kept in a 5% CO_2_ incubator set at 37°C.

### Cell culture carbon monoxide exposure

For treatment with CO, cells were transferred to a portable incubator (5% CO_2_, 37°C) set inside a fume hood. Compressed CO (200 ppm) was plumbed directly into the incubator. For the 5 hours of treatment, a constant flow of 5L/min at 1 psi CO was applied to the incubator. After the treatment, cells were returned to their home incubator.

### Nrf2 activity assay

Nrf2 activity was quantified using nuclear extracts and a TransAm NRF2 activity kit from Active Motif (cat# 50296) following the manufacturers included instructions.

### Cathepsin D activity assay

SH-SY5Y cells were grown on chamber slides to confluency. Cells were then treated with CO for five hours and returned to their home incubator. Twenty-four hours after starting CO treatment, active cathepsin D was stained for using the sir-Lysosome probe per the manufacturer instructions (Spirochrome, Rhein, Switzerland). Images were taken at 63X under oil immersion. Sir-lysosome fluorescence was measured by ImageJ and at least 15 cells were counted per condition.

### Empty Basket Quantification

Sections containing the cerebellum stained for calbindin and GAD were imaged at 40x using a VS120 scanning robotic microscope (Olympus). Empty baskets in the cerebellum were differentiated from full baskets by the presence of a GAD+ plexus of basket cell axons without a surrounded calbindin+ Purkinje cell body. Counting was restricted to the molecular-granule cell layer junction. Every cell within this junction was examined across three coronal sections of the cerebellum separated by at least 240 microns.

### Flouro-Jade C staining

Sections were stained for degenerating neurons using Fluoro-jade C (Biosensis cat# TR-100-FJ) per the manufacturer’s instructions. Sections were imaged at 40x using a VS120 scanning robotic microscope (Olympus). A total of 6 sections were counted per animal, each separated by at least 160 microns.

### Statistical analyses

Data are expressed as means ± SEM. Unpaired t-tests were used to analyze two groups. Unless otherwise stated, analysis of variance (ANOVA) was used for multiple comparisons, with Tukey post-hoc P values for pairwise comparisons. Pearson correlations were used. P < 0.05 was considered significant. Analyses were performed with GraphPad Prism.

## Abbreviations

PD: Parkinson’s disease
CO: carbon monoxide
AAV: adeno-associated virus
αSyn: alpha-synuclein
DA/TH^+^: dopamine/tyrosine hydroxylase-positive
MPTP: 1-methyl-4-phenyl-1,2,3,6-tetrahydropyridine
HO-1: heme oxygenase-1

## ACKNOWLEDGEMENTS

The authors thank Daniel Kalman (Emory, Department of Pathology) and Bradley Hyman (MGH, Department of Neurology) for insightful discussions, Joseph Locascio for statistical input, and Nitsan Goldstein (MGH, Department of Neurology) for research assistance. The authors thank Hillhurst Biopharmaceuticals for providing HBI-002 for these studies.

## Author Contributions

KNR contributed to the design, acquisition, analysis, and interpretation of data, as well as to the drafting of the manuscript.

MZ contributed to the design, to the acquisition, analysis, and interpretation of data, and to the review of the manuscript.

XX contributed to the acquisition, analysis, and interpretation of data, and to the review of the manuscript.

AF contributed to the acquisition, analysis, and interpretation of data, and to the review of the manuscript.

CW contributed to the acquisition, analysis, and interpretation of data, and to the review of the manuscript.

SL contributed to the design, to the acquisition, analysis, and interpretation of data, and to the review of the manuscript.

PW contributed to acquisition and interpretation of data and the review of the manuscript.

MAS contributed to interpretation of data and the review of the manuscript.

XC contributed to the design, analysis, interpretation of data, and the review of the manuscript.

SNG contributed to the conception, design, analysis, and interpretation of data, and to the drafting of the manuscript.

## Data availability

All data needed to evaluate the conclusions in the paper are present in the paper and/or the Supplementary Materials. The datasets generated and analyzed in the current study will be archived in Dryad. Requests for the HBI-002 reagent provided by Hillhurst Biopharmaceuticals should be submitted to: hillhurstbio.com/contact/.

## Materials and correspondence

Stephen Gomperts, MD, PhD

Massachusetts General Hospital Institute for Neurodegenerative Diseases

114 16^th^ Street

Charlestown, MA 02129

