## Supplementary data for "Neuroprotection of low dose carbon monoxide in Parkinson’s disease models commensurate with the reduced risk of Parkinson’s among smokers"

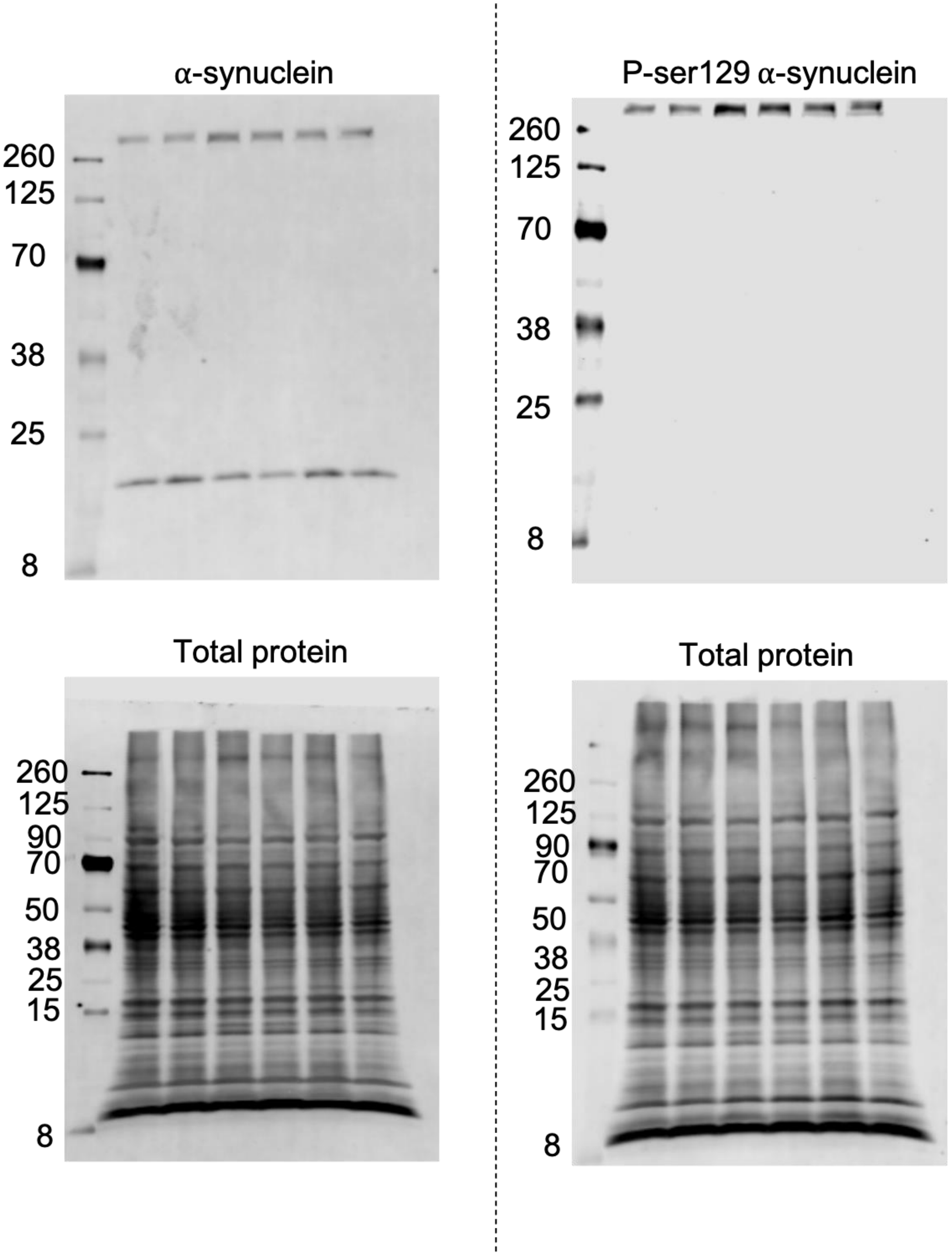


Supplementary Figure 1. Uncropped western blots of αSyn and p-ser129 αSyn

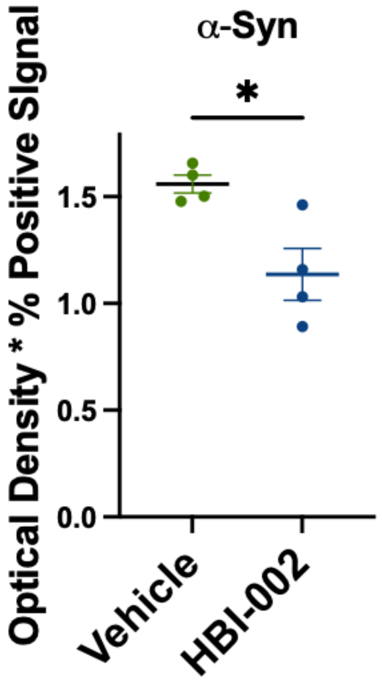


Supplementary Figure 2. Quantification of αSyn immunoreactivity in the substantia nigra pars compact of rats treated with vehicle or HBI-002. P=0.016, N=8, 4 per group.

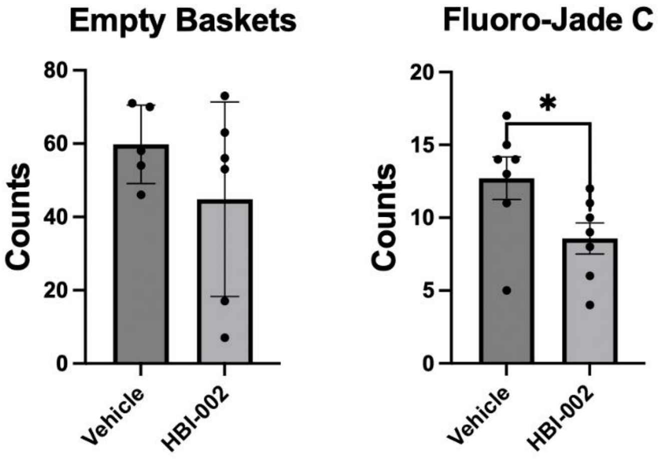


Supplementary Figure 3. Assessment of neurotoxicity. (**a**) Administration of HBI-002 had no effect on the number of empty basket counts (N=11, P=0.27). (**b**) HBI-002 reduced the number of Fluoro-Jade C positive cells in the prefrontal cortex compared to vehicle treated animals (N=14, P=0.041).

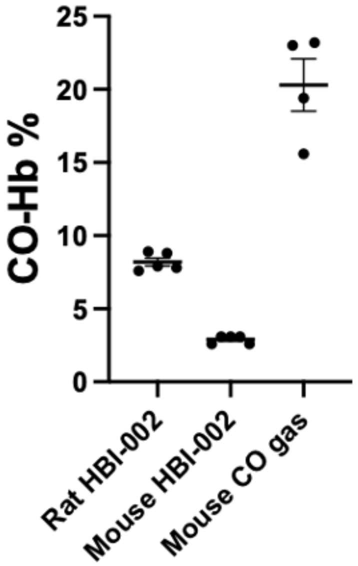


Supplementary Figure 4. CO route of administration and species effects on CO-Hb levels. HBI-002 10 ml/kg. CO gas, 200 ppm.

LC3b I-II

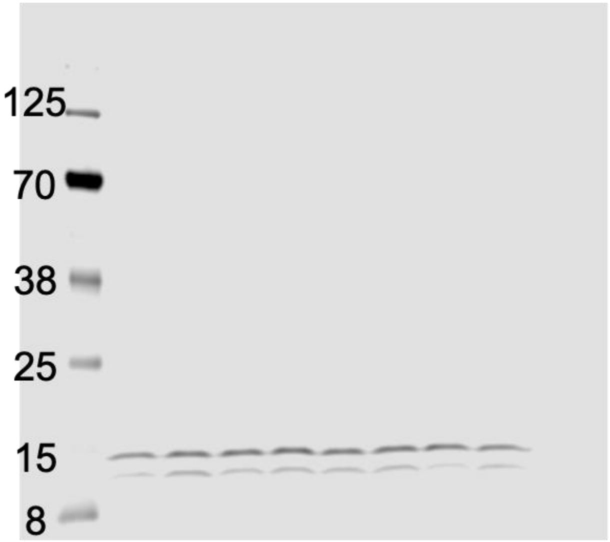


Total Protein

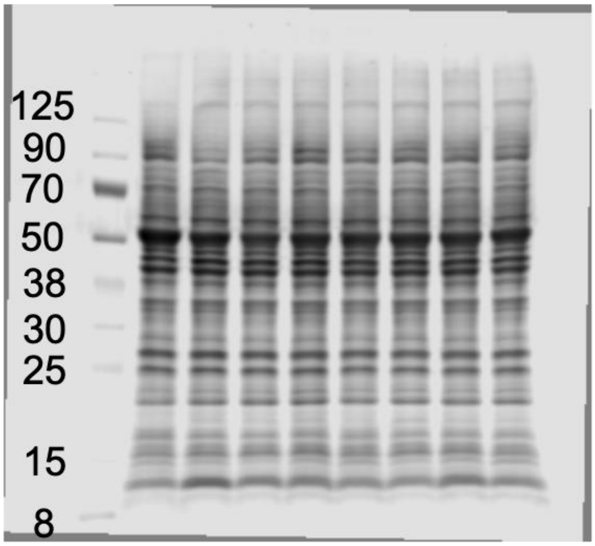


P62

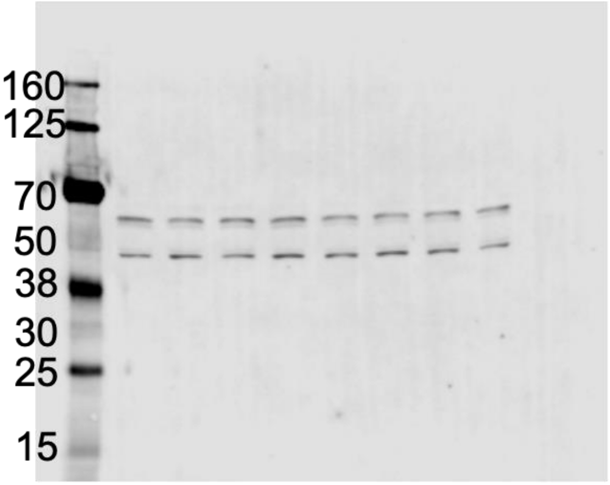


Total Protein

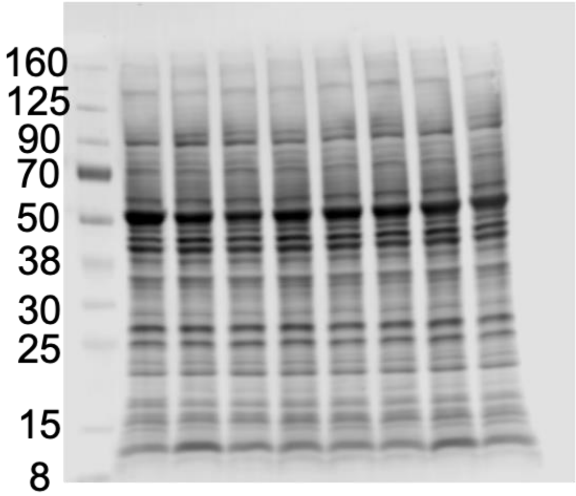

Rab4

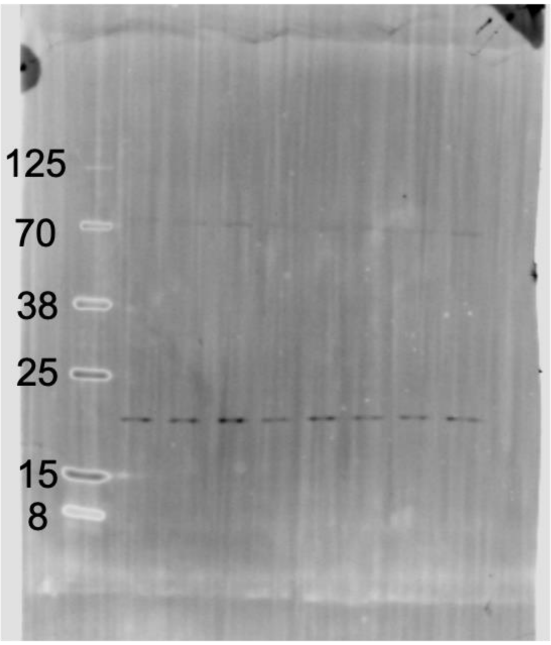


Total protein

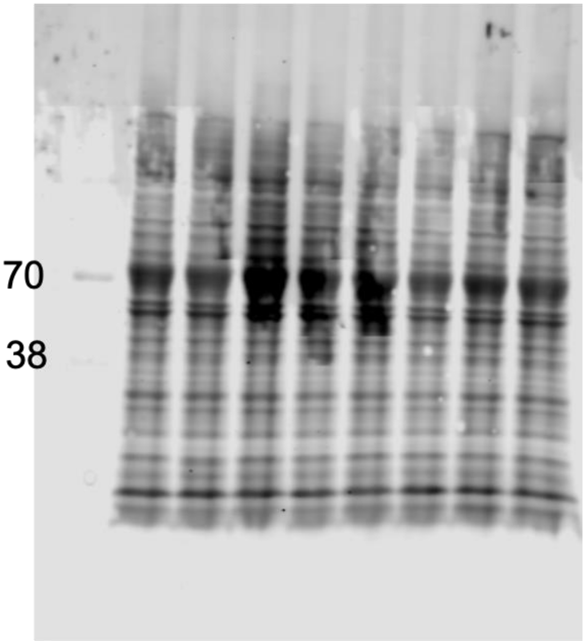


VSP35

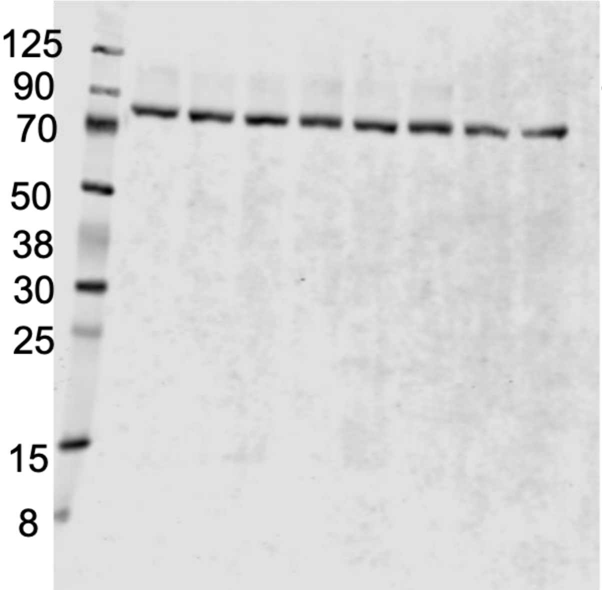


Total protein

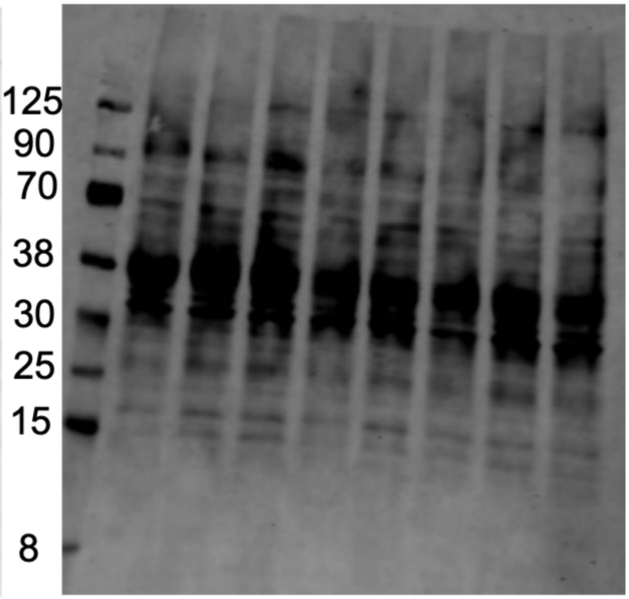

Lamp-1

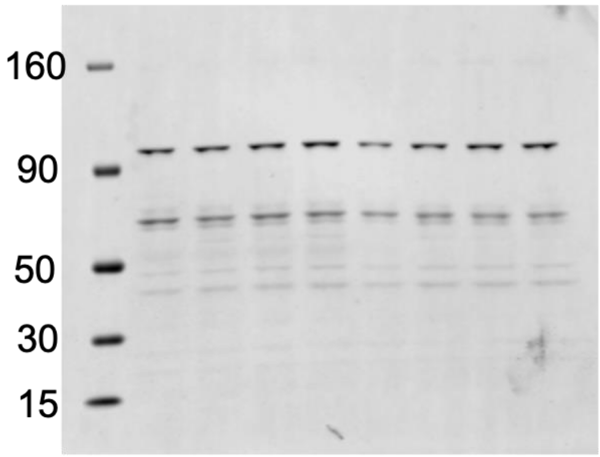


Total Protein

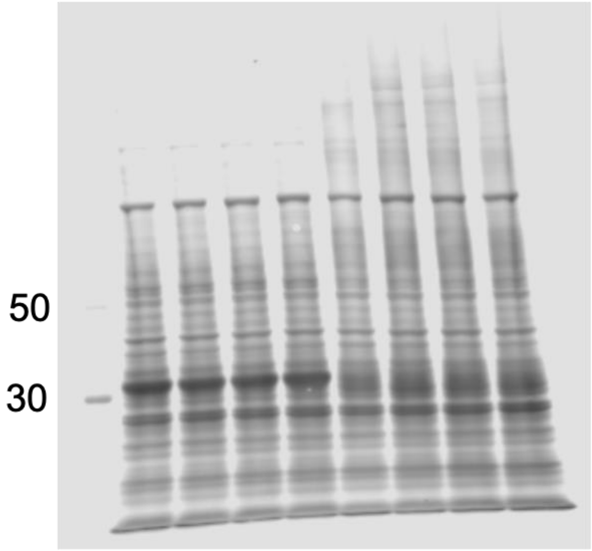


Ctsd

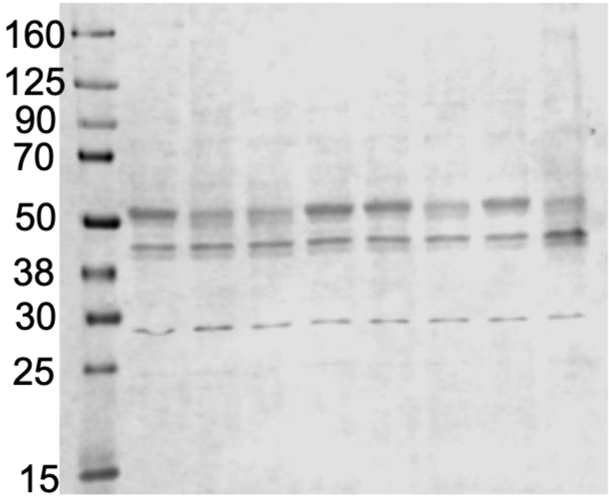


Total Protein

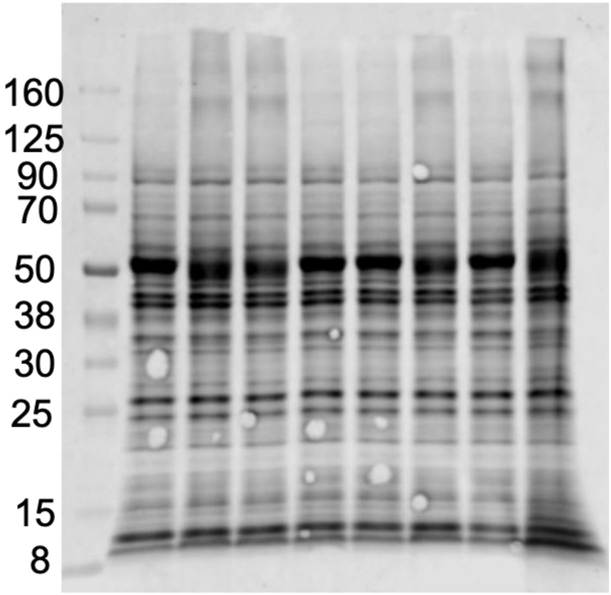

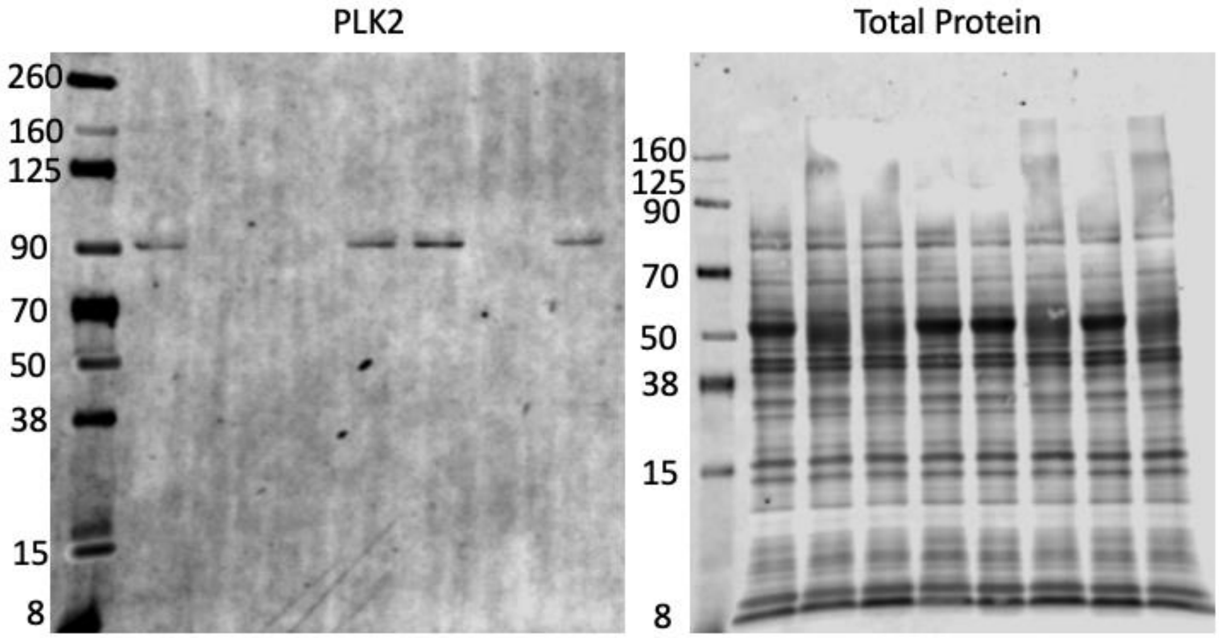


Supplementary Figure 5. Uncropped western blots for blots shown and quantified in Figure 2.

Hif-1α (Nuclear) Total protein

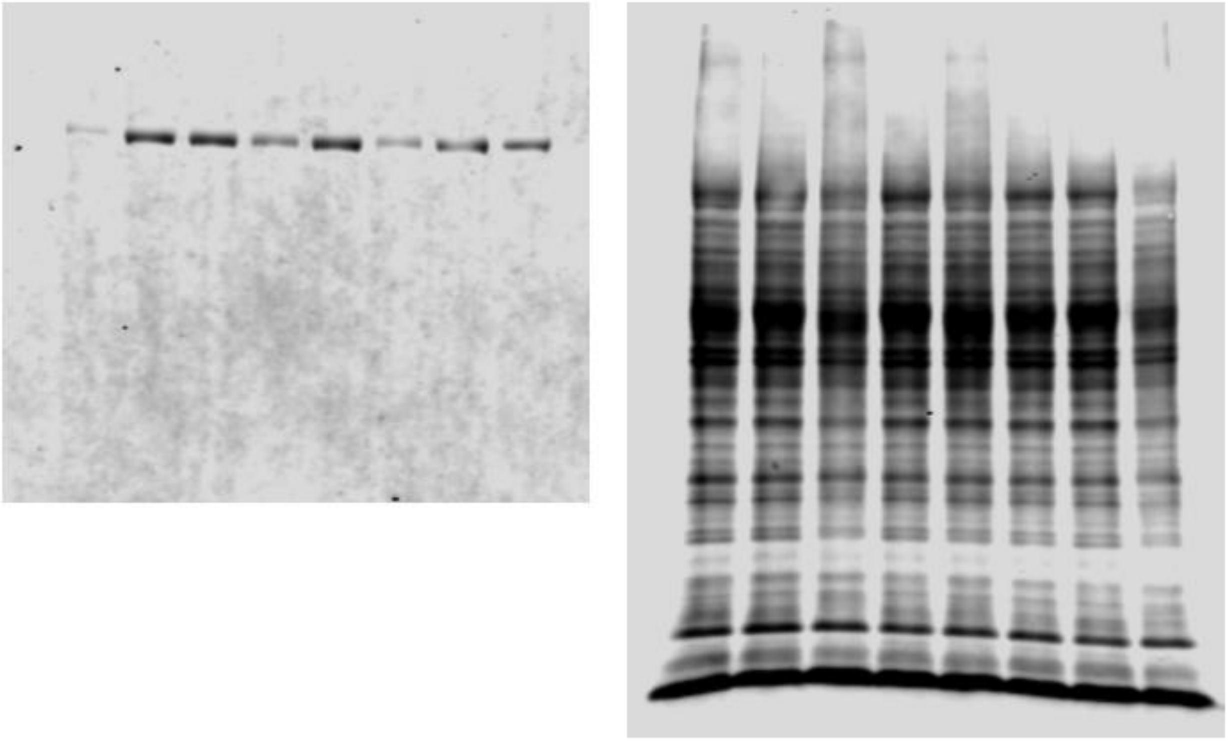

Supplementary Figure 6. Uncropped western blot for blot shown and quantified in figure 5.
